# Exploring the host range for genetic transfer of magnetic organelle biosynthesis

**DOI:** 10.1101/2023.01.31.526216

**Authors:** M.V. Dziuba, F.D. Müller, M. Pósfai, D. Schüler

## Abstract

Magnetosomes produced by magnetotactic bacteria have great potential for application in biotechnology and medicine due to their unique physicochemical properties and high biocompatibility. Recent studies uncovered the genetic determinants for magnetosome formation and inspired ideas to transfer the pathway into hitherto non-magnetic organisms. However, previous efforts to genetically magnetize other species have been successful in only a few bacterial recipients, revealing significant challenges in this approach. Here, by systematic examination of 25 proteobacterial hosts, we generated 7 novel magnetosome-producing strains. We further characterized the recombinant magnetosomes produced by these strains and delineate a set of auxiliary factors linked to magnetosome formation. Our findings will have significant implications for generation of magnetized life cells and the potential to facilitate the production of biogenic magnetic nanoparticles for a wide range of biomedical applications.

## Introduction

The idea to genetically endow magnetism to living cells has been of great interest in synthetic biology. For instance, magnetized microbes could be transformed into self-propelling and self-powered microrobots steered to deliver cargoes in microfluidic systems, or even in the human body^1–3^. By magnetic labelling, pathogens could be easily detected and recovered from infected hosts in laboratory studies^4^. Magnetic nanoparticles (MNP), produced by microorganisms or even higher eukaryotic cells may provide perfect contrast for magnetic imaging^5–7^. Magnetic actuation and radio wave heating of MNP were also suggested to remotely trigger gene expression and cellular functions in the emerging field of magnetogenetics^8–10^.

Biomagnetism is innate to magnetotactic bacteria, mud-dwelling microbes, which biomineralize chains of magnetic nanocrystals, so-called magnetosomes, with the exceptional properties for geomagnetic field sensing^11^. They are biosynthesized within the dedicated vesicles of the magnetosome membrane, in which physicochemical conditions are strictly controlled for the biomineralization of uniformly composed and chemically pure magnetite (Fe3O4) crystals with favorable narrow size distribution that cannot yet be replicated by chemical synthesis^12^. These unique properties, in combination with their biocompatibility and recent advances in functionalization through genetic engineering^13–15^ endorsed applications of MTB and magnetosomes in several biomedical fields, such as magnetic drug targeting, magnetic imaging, and hyperthermia cancer therapy^11,16–18^.

The unprecedented characteristics of magnetosomes also prompted ideas to genetically transfer their biosynthesis to other organisms. However, owing to the complexity of the process and an incomplete knowledge of its genetic determinants, this has been accomplished only recently, and only in a few prokaryotic recipients. In the widely studied *Alphaproteobateria* MTB *Magnetospirillum gryphiswaldense* and its close relatives, clusters of about 30 *mam* and *mms* genes were identified to control magnetosome biosynthesis, which have specific functions in iron transport, magnetosome membrane formation, and crystal biomineralization, as well as organizing and positioning of magnetosome chains. Transfer of this gene set induced biosynthesis of well-ordered magnetosome chains in the photosynthetic bacterium *Rhodospirillum rubrum*, demonstrating that the magnetic organelle can be fully reconstituted within foreign recipients by balanced expression of structural and catalytic factors. Later, the closely related but non-magnetotactic chemotrophic *Magnetospirillum* sp. 15-1^19^ and a photosynthetic *Acetobacteraceae* species *Rhodovastum atsumiense* G2-11 were also shown to synthesize small imperfect magnetosomes upon transfer of the same set of genes, although the trait proved highly unstable in the latter^20^.

These findings have stimulated enormous efforts within the synthetic biology community to transfer magnetosome biosynthesis into more facile hosts, and even inspired ideas to borrow genetic parts from MTB for endogenous magnetization of cells from higher organisms ^21–23^. Heterologous magnetosome biosynthesis in other, biotechnologically relevant and more tractable hosts would also alleviate some problems associated with the challenging cultivation and limited tractability of the available native and synthetic magnetic bacterial strains^24,25^. However, despite the encouraging results, simple transfer of magnetosome genes has failed for several other potential hosts. This includes numerous futile efforts by several labs^26^, including ours (Dziuba et al., unpublished) to reconstitute magnetosome biosynthesis in the most prominent biotechnological model and production organism *E. coli*, and is also illustrated by the regular attempts to genetically magnetize *E. coli* in the well-known international *Genetically Engineered Machine competition* (iGEM)^27–29^. While the exact reasons for these failures are unknown, it can be speculated that besides *mam* and *mms* gene clusters, some additional regulatory functions, precursors, and co-factors are required for magnetosome formation and might be present in some but absent from other putative hosts. Thus, the gaps in our understanding of magnetosome biosynthesis pose a significant challenge for rational engineering of further organisms for this process, and successful hosts have been identified so far only by chance.

Here, we tested a range of various bacteria for the ability to produce magnetosomes by attempting the transfer of the canonical set of magnetosome genes from *M. gryphiswaldense* to 25 different proteobacterial species (Fig. 1). This allowed us to successfully transform 14 strains and generate 7 novel transgenic strains able to stably produce magnetosomes, including several that are based on species broadly used in biotechnology. We further characterize the recombinant magnetosomes produced by these synthetic strains and delineate a set of auxiliary factors potentially linked to magnetosome formation by comparing the genomes of the magnetosome-competent (Mag+) and magnetosome-incompetent (Mag-) species. Our findings will have significant implications for the generation of magnetized organisms and the potential to facilitate the production of biogenic magnetic nanoparticles for a wide range of biomedical applications.

**Fig. 1.**
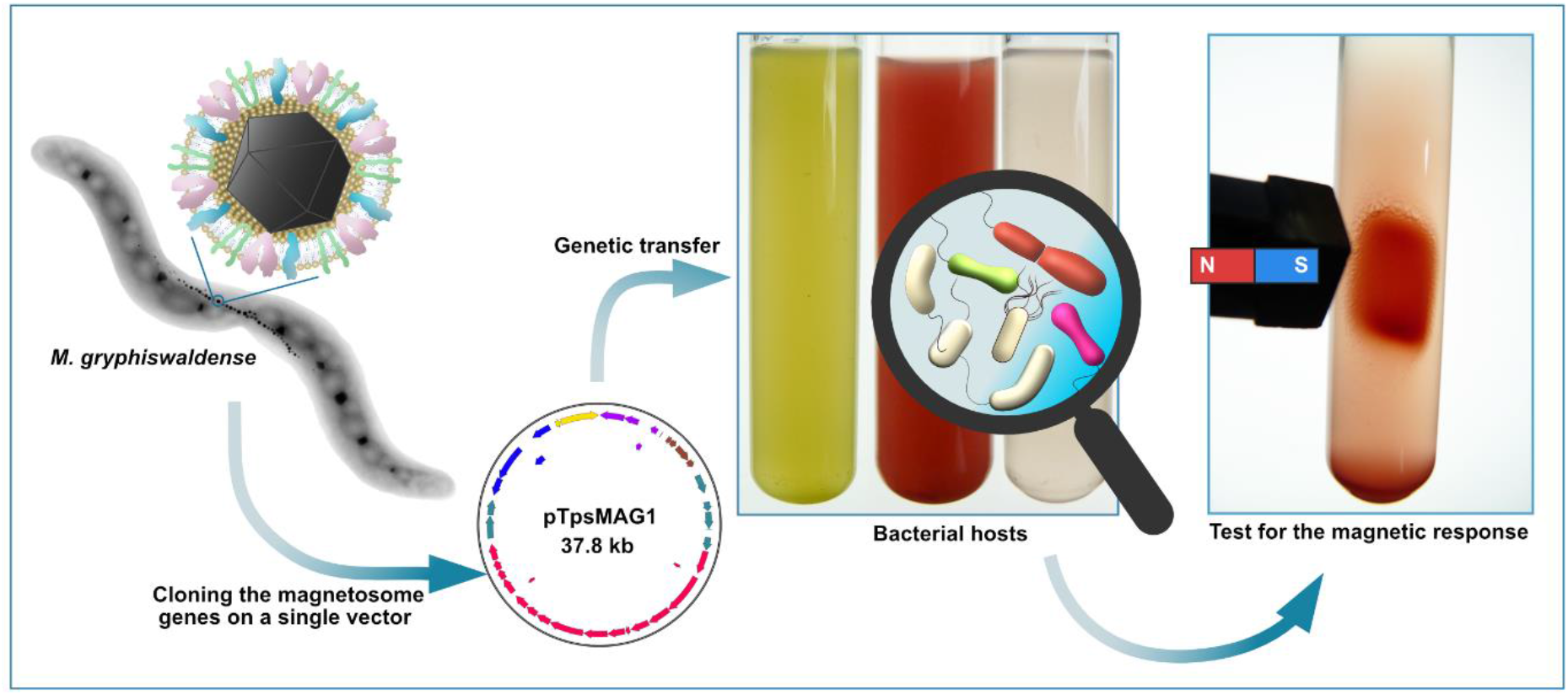
Experimental strategy of the survey for bacterial hosts for heterologous magnetosome production. The genes encoding magnetosome biosynthesis pathway from M. gryphiswaldense were cloned on a single compact vector pTpsMAG1 that was used to transform 25 bacterial strains belonging to different phylogenetic groups within Proteobacteria.

## Results

The five major magnetosome gene operons *mamAB, mamGFDC, mms6op, mamYXZ*, and *feoABm from M. gryphiswaldense* were delivered by conjugation of the single plasmid pTpsMAG1^19^, which becomes randomly integrated into the genome of the target hosts by means of the mariner transposon (Fig. 1). Integration was successful in 14 out of 25 strains tested (in the following referred to as MAG transconjugants), whereas in 11 species no insertions could be detected despite repetitive attempts (Fig. S1, Table S1). In some of the untransformed strains, e.g., *P. photometricum, M. fulvum*, or *M. molischianum*, this failure can be attributed to the unavailability of reliable transformation methods, while in others, the potential high metabolic burden or toxicity of the magnetosome genes expression could lead to negative selection against the insertion^30^. Conspicuously, in one of the successfully transformed strains, *Rh. centenaria*, a deletion eliminating most of the magnetosome gene cassette was repeatedly detected in all screened transconjugants from several independent transformation experiments (Fig. S2).

However, transfer of the gene set induced the synthesis of magnetosome-like particles in *Cereibacter sphaeroides, Rhodoplanes elegans, Rhodopseudomonas pseudopalustris, Blastochloris viridis, Rhodoblastus acidophilus, Azospirillum brasilense*, and *Rhodomicrobium vannielii* (Fig. 2, hereafter referred to as Mag+). In 5 recipient strains, no magnetosomes were detected: *Agrobacterium tumefaciens, Mesorhizobium japonicum, Rhodobacter capsulatus* SB1003, *Rba. capsulatus* B10, and *Rubrivivax gelatinosus* (Fig. S3, Mag-), although the magnetosome cassette in these transconjugants remained intact as confirmed by both PCR and whole genome re-sequencing (not shown).

**Fig. 2.**
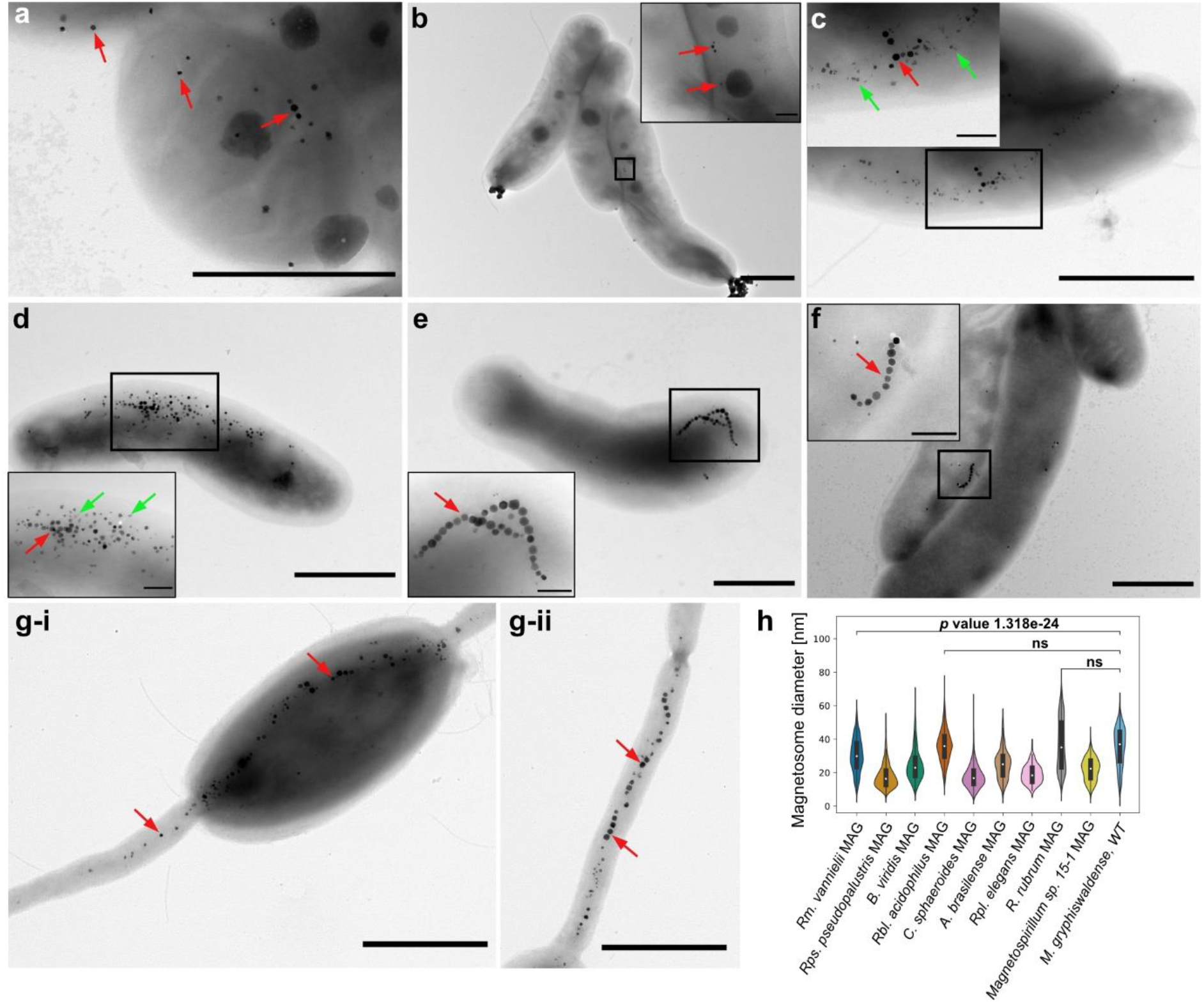
Overview of the magnetosome producing hosts. (a - g) Electron micrographs of (a) C. sphaeroides MAG, (b) Rpl. elegans MAG, (c) Rps. pseudopalustris MAG, (d) B. viridis MAG, (e) Rbl. acidophilus MAG, (f) A. brasilense MAG, (g) Rm. vannielii MAG with the magnetosome chain localized mostly in the cell body (g-i) and in the prostheca (g-ii). (h) Violin plot showing size distribution of the magnetosomes produced by the foreign and native (M. gryphiswladense) hosts. Scale bars: 1 μm. ns: not significant.

Noteworthy, all but one transgenic strain found capable of magnetosome biosynthesis in this study grow preferably photoheterotrophically under anaerobic conditions, and this growth regime apparently stimulated magnetosome formation in comparison to microoxic chemotrophic cultivation as also demonstrated previously for *R. rubrum*^25,31^. The strains synthesizing more numerous magnetosomes, e.g., *Rm. vannielii* MAG (Supplementary video V1)*, Rbl. acidophilus* MAG (magnetically attracted culture in Fig. 1), and *B. viridis* MAG (Fig. S4), demonstrated conspicuous magnetic response and could be attracted by a bar magnet. Analysis of the magnetosomes by STEM-EDS imaging revealed that all of them were composed of iron (Fe) and oxygen (O) with traces of phosphorus (P), likely originating from the magnetosome membrane (Fig. 3-4, Extended Data Figures 1-2).

**Fig. 3.**
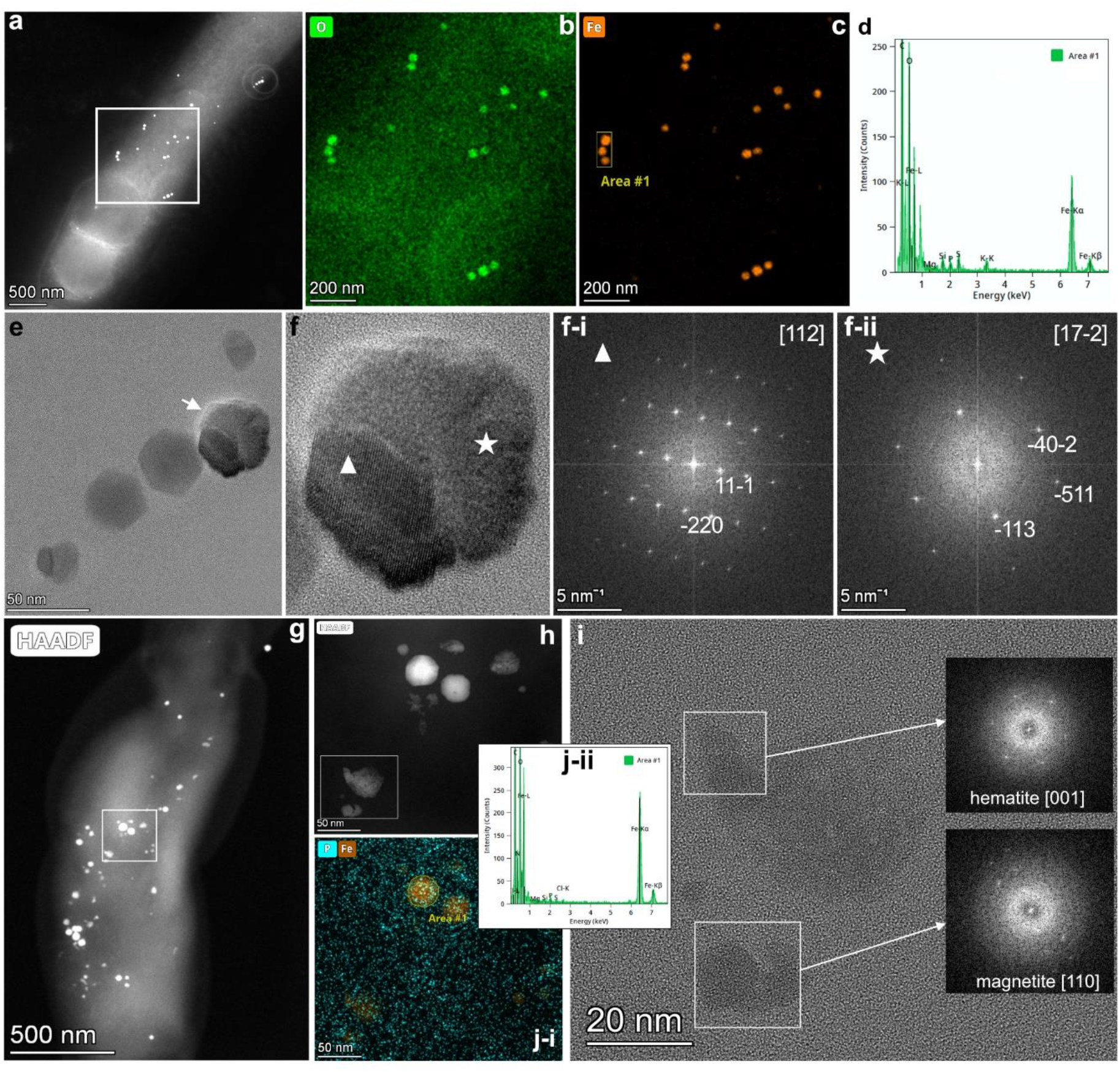
*Crystallographic analysis of the magnetosomes produced by C. sphaeroides* MAG (a-f) and *Rps. pseudopalustris* MAG (g-j). (a) HAADF of a *C. sphaeroides* MAG cell; (b) and (c) elemental maps of the boxed area in (a). (d) The EDS spectrum of Area #1 in (c). (e) HRTEM of the magnetosomes from *C. sphaeroides* MAG. (f) HRTEM of a crystal marked in (e) with arrow. (f-i) FFT of the left (marked with a triangle); (f-ii) FFT of the right (marked with a star) intergrown crystals, both consistent with magnetite, in [112] and [17-2] orientations, respectively. (*g) HAADF of a Rps. pseudopalustris* MAG cell. (h) close-up of the area highlighted with the white rectangle in (g). (i) HRTEM image of the flake-shaped magnetosomes highlighted with the rectangle in (h) with FFT of the boxed areas. The magnetite particle is in the [110] zone axis orientation, the hematite particle is in close to [001] orientation. (j-i) Elemental map of the close-up from (h) and (j-ii) EDS spectrum of Area #1 in (j-i).

**Fig. 4.**
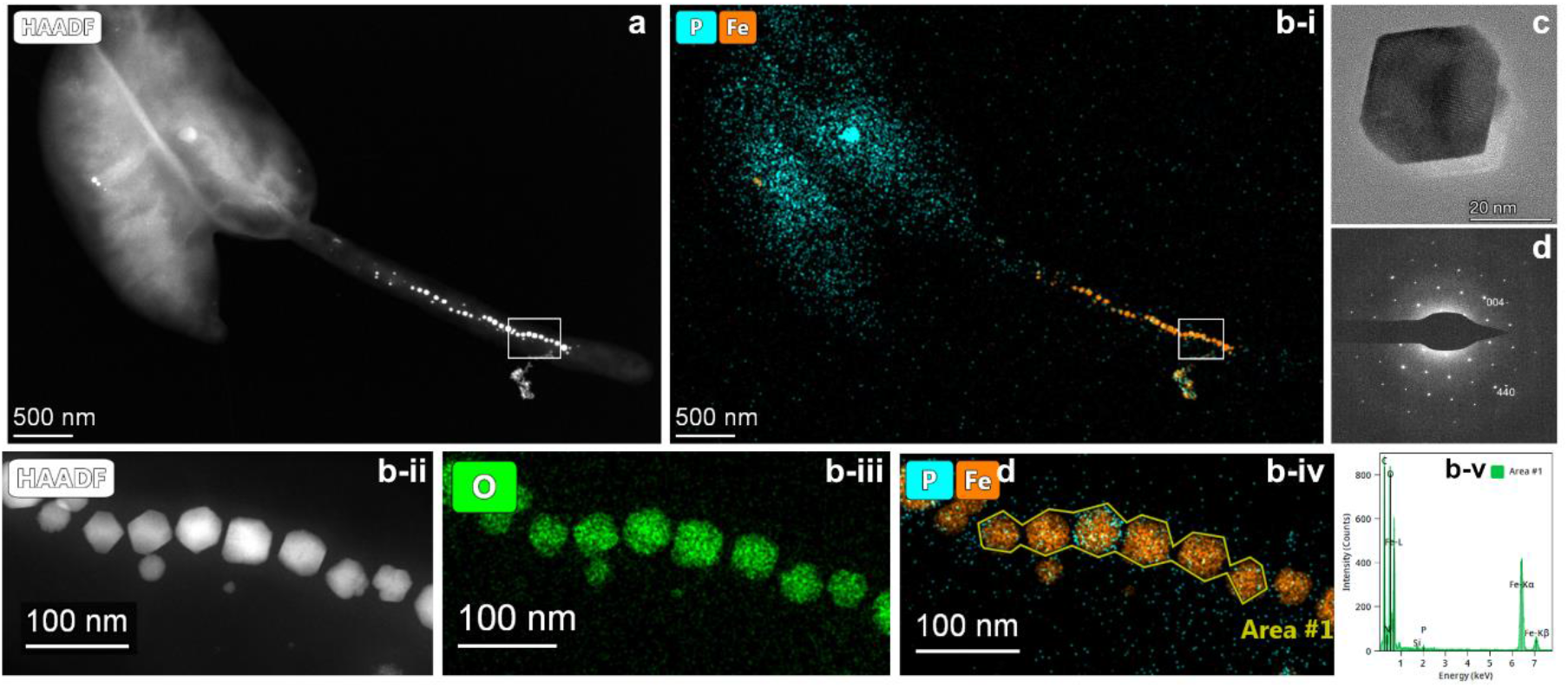
HAADF image (a) and elemental maps (P, Fe and O) of representative Rm. vannielii MAG cells (b), showing a magnetosome chain in the prostheca. (b-ii – b-iv) Close-up HAADF and elemental maps of the chain fragment highlighted with the white rectangle in b-i, (b-iv) EDS spectrum of Area #1 in b-iv. (c) HRTEM image of a typical magnetite crystal from Rm. vannielii MAG with (d) a SAED pattern in [110] orientation.

We failed to detect any magnetosome-like particles in *C. sphaeroides* MAG under its previously reported optimal growth conditions in Sistrom medium^32^ supplemented with 50 μM ferric citrate, but when cultivated in the standard medium typically used for *M. gryphiswaldense* (FSM), cells synthesized small particles (17.70 ± 6.63 nm), scattered throughout the cells, or occasionally arranged in short chains (Fig. 2a, h, Fig. 3a-e). HRTEM revealed that most of magnetosomes in *C. sphaeroides* contained twinned or tripled magnetite crystals (Fig. 3f), compared to the single crystals predominating in the donor MTB *M. gryphiswaldense*. In contrast, *Rpl. elegans* demonstrated only rudimentary magnetosome biosynthesis by forming 1-3 tiny (19.19 ± 5.33 nm) magnetosomes per cell (Fig. 2b, h).

*Rps. pseudopalustris* MAG formed short chains or agglomerates comprising 3-5 large magnetosomes composed of magnetite, accompanied by numerous scattered, small, aberrant, flake-shaped particles containing hematite and other iron oxide crystals (Fig. 2c; Fig. 3g-i). Altogether, the magnetosomes in *Rps. pseudopalustris* MAG were numerous (42.23 ± 27.67 magnetosomes/cell) with the average size of 17.81 ± 6.97 nm (Fig. 2h). The appearance and composition of the magnetosomes synthesized by *B. viridis* MAG were similar to those observed in *R. palustris* MAG; however, the large magnetite-containing magnetosomes were more numerous, whereas flake particles were much less abundant (Fig. 2d). This resulted in a larger average crystal size (24.60 ± 9.29 nm) than in *R. pseudopalustris* MAG (Extended Data Fig. 1). Although the magnetosomes from *B. viridis* MAG were composed of pure magnetite, the crystals were often irregularly shaped, e.g., twinned or intergrown (Extended Data Fig. 1).

*Rbl. acidophilus* differs from the other strains used in this study by its preferred growth under slightly acidic conditions^33^. Although a low pH (pH 6.0) was shown to severely impair magnetite crystal biomineralization in *M. gryphiswaldense*^34^, Rbl. acidophilus MAG cultivated in the acidic medium (pH 5.5), that is optimal for growth, formed extended magnetosome chains, along with some scattered and clustered magnetite particles (Fig. 2e). However, many crystals had aberrant shapes reminiscent of those formed at low pH by *M. gryphiswaldense*^34^ (Extended Data Fig. 2). When grown in media with various pH (5.5, 6.0, 6.5, and 7.0), *Rbl. acidophilus* MAG formed the longest chains consisting of perfectly shaped magnetosomes at pH 7.0, despite the growth was considerably diminished (Extended Data Fig. 3). Of note, the strain naturally forms intracellular iron- and phosphorous-rich granules, which are compositionally close to ferrosomes, the recently discovered biomineral-containing organelles formed by some proteobacteria^35^ (Extended Data Fig. 2b-e, g).

Among the strains tested in this study, *A. brasilense* was the only non-phototrophic species able to produce magnetosomes. However, microoxic conditions that are known to support magnetosome biosynthesis in the gene donor *M. gryphiswaldense*, entirely repressed magnetosome biosynthesis in this strain. At the same time, under the denitrifying conditions, i.e., anoxic in the presence of the reducing agent L-cysteine and increased nitrate concentrations (12-20 mM), relatively small, scattered magnetosomes were formed that occasionally assembled into chains (Fig. 2f, h).

Of all transgenic strains generated, the prosthecate *Hyphomicrobiales* species *Rm. vannielii* produced magnetosomes that were the most similar to those in *M. gryphiswaldense* in both morphology and intracellular chain organization, with the predominance of well-formed cuboctahedral magnetite crystals with an average size of 30.44 ± 9.95 nm (Fig. 2g, h; Fig. 4). However, due to the unique morphology of the strain, the magnetosome positioning was unusual: the chains often spanned across the cell body, and even extended towards, or occurred exclusively within the prosthecae (Fig. 4a, b).

Next, we asked which auxiliary factors might be distinctive for successful magnetosome formation by Mag+ foreign hosts, compared to Mag-. Previous genetic analyses^36–42^ had established that some genes located outside magnetosome clusters are also involved in magnetosome biosynthesis in *M. gryphiswaldense* (hereafter referred to as **<u>au</u>**xiliary **<u>g</u>**enes, **AUGs**). Although none of these genes appeared to be absolutely essential for magnetosome formation in *M. gryphiswaldense*, we hypothesized that the absence of certain genes might result in the inability of a foreign organism to produce magnetosomes. Therefore, we first screened all the Mag+ and Mag- hosts, including the previously reported^19,20,43^, for the presence of AUGs (see Materials and Methods for details). Of 90 AUGs, predicted in the previous studies, only 25 were found present in the genomes of all Mag+ (in the following, **<u>f</u>**iltered **<u>a</u>**uxiliary **<u>g</u>**enes, **FAG**s). However, among FAGs there were none that would be absent in ALL Mag-, suggesting that the lack of the desired phenotype in them cannot be simply attributed to the universal absence of a certain gene(s) from the currently known AUGs set (Fig. 5a). Nonetheless, we could not rule out that the lack of different AUGs can account for the failure to produce magnetosomes in each individual host. For example, a closer inspection of the AUGs’ distribution with the experimentally confirmed link to magnetosome formation revealed that *Rvx. gelatinosus* lacks the genes encoding DsbAB-like proteins potentially involved in disulphide bond formation, inactivation of which was shown to severely affect magnetosome biosynthesis in *M. gryphiswaldense*^36^ (Fig. 5b). However, we could not find any further patterns in the AUGs’ distribution that would correlate with the ability to synthesize magnetosomes, or at least flake crystals, as in *Rps. pseudopalustris* MAG and *B. viridis* MAG.

**Fig. 5.**
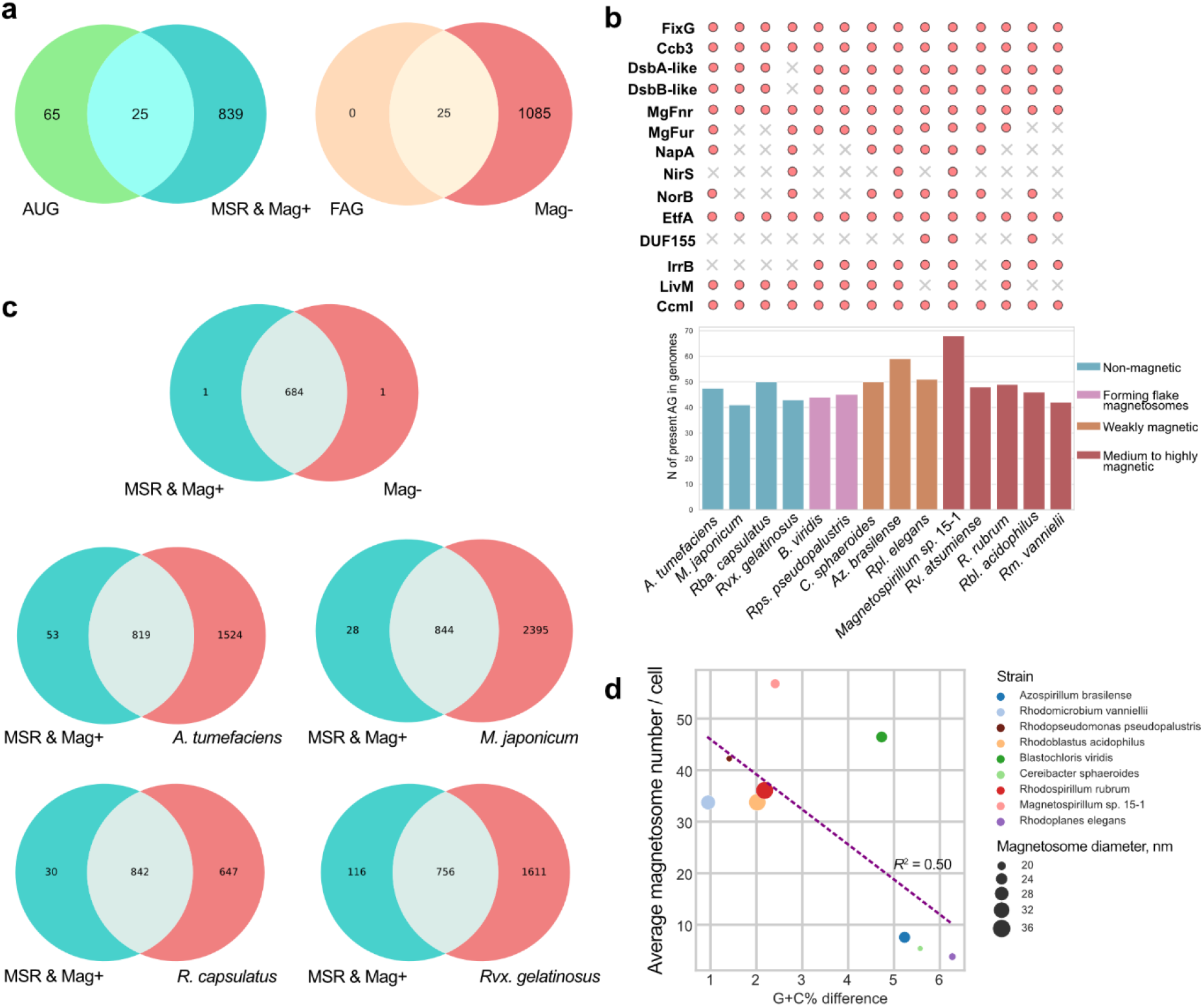
Genome-genome comparisons of Mag+ and Mag- strains. (a) Venn diagrams representing (left) AG shared by M. gryphiswaldense (MSR) and Mag+, and (right) which of the AG shared by Mag+ are also shared by Mag-. (b) Bar chart demonstrating the number of AG present in all strains used in this study and those previously magnetized. The upper diagram shows distribution of the AG with experimentally confirmed phenotypes (circle: gene present, cross: gene absent). (c) Venn diagrams representing the fractions of orthologues shared between the combined MSR & Mag- group and combined of individual Mag- strains. SB1003 was used as representative strain for R. capsulatus in this analysis. (d) Relational plot demonstrating moderate negative correlation (Pearson’s r −0.707, p value 0.033) between the magnetosome number per cell and the difference in the GC-content between M. gryphiswaldense and the host’s genome (as modulo). The symbol size indicates the average magnetosome diameter.

Since the full set of AUGs is still undefined, we could not completely rule out that Mag- strains might be lacking potentially important genes, for which linkage to magnetosome biosynthesis has not yet been established. Therefore, we further compared all the genes present in the hosts in the all-against-all approach. Comparison of the orthologues set shared by Mag+ strains and *M. gryphiswaldense* against Mag- strains revealed only a single orthologue (*cheW*) found in all Mag+ but absent in Mag- (Fig. 5c). However, the encoded chemotaxis protein CheW was shown to be important for aerotaxis but not for magnetosome biosynthesis in *M. gryphiswaldense*^44^, and is also unlikely to be associated with magnetosome biosynthesis in the other hosts.

Further exploration revealed several candidate genes, which, based on their predicted function, might contribute to magnetosome biogenesis. For example, both *A. tumefaciens* and *M. japonicum* lack the ferrous transport system FeoAB that is homologous to FeoAB2 in *M. gryphiswaldense*. The latter was shown to contribute to magnetosome formation, although to a much lesser extent than FeoAB1 (FeoABm)^45,46^, which was part of the magnetosome gene set transferred to all hosts on TpsMAG1. Hence, the deficiency in ferrous transporters is not a likely explanation for the absence of magnetosomes. Besides, the Mag+ group shares various genes encoding proteins with unknown functions that are not found in one or several Mag- strains, and hence their contribution to magnetosome formation cannot be excluded.

Apart from the presence of potential AUGs, we observed that all Mag+ hosts share a common metabolic characteristic: they are capable of anaerobic growth, supported either by denitrification (*Magnetospirillum* sp. 15-1 and *A. brasilense*) or photosynthesis. A link between denitrification and magnetosome formation is well established in *M. gryphiswaldense*, where it was suggested to play a role in the oxidation of ferrous iron supporting magnetite biomineralization under both anoxic and microoxic conditions^38,47^. Compared to *M. gryphiswaldense*, in which nitrate respiration overlaps with oxygen respiration under microoxic conditions, activation of denitrification in *A. brasilense* requires much lower oxygen concentrations^48^. Therefore, our observation that magnetosome production in *A. brasilense* MAG became initiated only under anoxic conditions reinforces the significance of denitrification for magnetosome biogenesis. Consistent with this, the chemotrophic Mag- strains have only incomplete denitrification pathways (*A. tumefaciens*) or are entirely devoid of this pathway (*M. japonicum*), which is also reflected by the absence of some or all of the associated genes in the AUGs’ distribution pattern (*napA, nirS, norB*, Fig. 5b). Besides, *A. tumefaciens* reduces nitrite using NirK instead of NirS, which is not homologous to NirS and might be unable to replace it in magnetosome biosynthesis. We also noticed that anaerobic growth of *A. tumefaciens* with nitrate in batch cultures was very slow in comparison to aerobic cultivation, which could undermine magnetosome biosynthesis due to energy limitations. This can be explained by unbalanced expression of the denitrification enzymes occurring frequently during the aerobic-anoxic transition and resulting in uncontrolled accumulation of NO to toxic levels observed in this strain previously^49,50^. All these factors might have contributed to the lack of magnetosome biomineralization in *A. tumefaciens*.

The potential link between magnetosome biogenesis and photosynthesis is more enigmatic. To date, no phototrophic magnetotactic bacteria have been found in nature. Even in the phototrophic *Rhodovastum atsumiense*, in which a complete set of silent magnetosome genes was discovered recently, does not produce magnetosomes under lab conditions^20^. At the same time, the magnetosome production was clearly stimulated by phototrophic conditions in all the phototrophic strains generated in this and previous studies, whereas none of them has a complete denitrification pathway. Among the factors that are common for anoxygenic photosynthesis, nitrate respiration, and magnetosome biogenesis, the involvement of *c-*type cytochromes is the most conspicuous. It can be hypothesized that cellular *c-*type cytochromes are directly involved in the ferrous ion oxidation required for the magnetite biomineralization. Alternatively, biogenesis of *c-*type cytochromes for all three pathways requires activation of the cytochrome *c* maturation system Ccm, an implication of which in magnetosome biogenesis was revealed in Silva and Schüler et al. (2020)^36^. Future experimental work should untangle the intricate link between magnetosome biogenesis and these two pathways.

Alternatively, lack of as well as variations in the magnetic phenotype could be attributed to poor expression of the introduced magnetosome genes. Although we did not estimate the gene expression directly, we found that differences in the G+C% content between the Mag+ hosts and *M. gryphiswaldense* had moderate negative correlation with the number of magnetosomes produced per cell (Pearson’s r −0.707 / *R*^2^ = 0.50, *p* value 0.033, Fig. 5d). Since G+C% content is a major factor shaping codon usage^51^ and also influencing the stringency of the promoter’s canonical σ70 motif^52^, the expression of the magnetosome genes was likely more efficient in the hosts with the lower G+C% difference. While most of the Mag- had relatively close G+C% to *M. gryphiswaldense*, more phylogenetically distant *Rvx. gelatinosus* showed the highest G+C% difference among all the hosts (7.7%, not shown on the plot). Therefore, at least in this host, the lack of magnetosome production might be also attributed to the potentially low expression.

## Discussion

Here, we demonstrate that biosynthesis of magnetosomes can be transferred to a wide range of bacteria, including some that are more distantly related to the gene donor *M. gryphiswaldense* than previously magnetized bacteria. However, heterologous magnetosome production has remained so far limited to the representatives of *Alphaproteobacteria*, likely because the constraints imposed by differences in promoter recognition, codon usage, and metabolic backgrounds become more pronounced with the expanding phylogenetic distance.

Our results revealed that no single gene, or gene set could predict the ability to biosynthesize magnetosomes by foreign hosts upon the transfer of magnetosome genes, whereas lack of different AUGs might have contributed to the failure to produce magnetosomes in certain host strains. At the same time, we observed that the ability to perform either denitrification or photosynthesis is crucial for magnetosome biogenesis, at least with the pathway acquired from *M. gryphiswaldense*. On the one hand, this reinforces the previously suggested link between iron oxidation during magnetite biomineralization and electron transport chains^37,39^. On the other hand, magnetosome biosynthesis can also potentially co-utilize the cytochrome *c* biogenesis system that is activated for production of ample amounts of cytochromes required for either nitrate respiration or photosynthesis, for maturation of the magnetochromes^53,54^. However, none of these observations would explain the lack of magnetosomes in *Rba. capsulatus*, which in this species might result from some regulatory mechanisms suppressing expression of foreign genes. Further research will be needed to understand these mechanisms and to develop strategies for overcoming them. Nonetheless, our findings shed light on the metabolic pathways supporting magnetosome biogenesis and provide a framework for future studies that will enable magnetosome production even in the hosts initially lacking the permissive background by their metabolic engineering.

Overall, our study provides important insights into the potential and challenges of heterologous magnetosome biosynthesis and paves the way towards engineering more distant hosts by systematic elimination of these barriers. Moreover, the phototrophic lifestyle of most of the magnetosome-producing hosts makes them ideal for light-powered magnetosome biosynthesis, which can reduce the costs and enable more sustainable magnetosome production. Two of such strains, *C. sphaeroides* and *Rps. pseudopalustris*, are organisms of biotechnological importance that are being used, for example, to produce biogas and valuable secondary metabolites^55,56^. This makes them attractive as potential multiproduct platforms that can be engineered for simultaneous synthesis of magnetosomes along with other products to further enhance the cost efficiency of magnetosome fabrication. As *C. sphaeroides* was shown to synthesize a non-pyrogenic type of lipopolysaccharides^57,58^, further optimization of this strain for improved magnetosome biosynthesis could enable the production of endotoxin-free magnetosomes for potential biomedical applications. We envision that the collection of transgenic magnetosome-producing strains constructed and characterized in our study will be a valuable resource for the development of innovative diagnostic and therapeutic methods based on magnetized bacteria and magnetosomes.

## Methods

### Bacterial strains and cultivation conditions

Strains used in this study, their cultivation conditions, and antibiotic concentrations for the selection of transconjugants are indicated in Table S2. For colony formation, the corresponding media were supplemented with 1.5% (w/v) of agar. Phototrophic strains were illuminated using tungsten light bulbs with the intensity 50 μmol/s/m2. To provide anaerobic conditions, the strains were incubated in Hungate tubes filled with medium to 2/3 of the volume and the headspace filled with 100% nitrogen, without agitation. If colony formation required anaerobic conditions, agar plates were incubated in anaerobic jars filled with 100% nitrogen and suppled with Anaerocult™ A reagent packages (Merck KGaA, Darmstadt, Germany).

### Genetic transformation

The plasmid pTpsMAG1 was transferred by biparental conjugation using *E. coli* WM3064 (*thrB1004 pro thi rpsL hsdS lacZΔM15* RP4-1360Δ (*araBAD) 567ΔdapA1341::[erm pir]*) as donor, largely according to the protocol described elsewhere^59^. The donor strain was refreshed by diluting the overnight culture 1:20 in the LB medium supplied with DAP and appropriate antibiotics 3-4 hours before mixing with the recipient culture. The procedure varied between the strains in the applied antibiotic (kanamycin or spectinomycin) concentration and the conditions for incubation of agar plates, which are summarized in Table S2. The obtained transconjugants were PCR-screened for the presence of the integrated magnetosome genes using the primers and procedure published previously^19^.

### Genome sequencing and assembly

To check the obtained mutants for the integrity of the MAG cassette, genome sequencing was performed. To this end, genomic DNA was extracted from 2-10 mL of stationary cultures with the Zymo Research Midiprep gDNA kit (Zymo Research EUROPE GmbH, Germany). Library preparation and sequencing were performed at Novogene (UK) Co. Ltd using Illumina NovaSeq 6000 with paired-end 2 × 150 bp reads corresponding to 1.0-1.3 Gbp in different samples. For the raw reads produced with Illumina, adapter trimming and quality control filtering were carried out with *fastp* using the default parameters^60^. Processed reads were mapped to the reference genome using *Bowtie2* ^61^ and Geneious (8.1.9)^62^. Various mutations were detected only in the variable region of the non-essential gene *mamJ* in the magnetosome cassette in most of the analyzed mutants, which however did not lead to frameshifts and is also known to not affect magnetosome formation. Otherwise, no mutations have been detected.

### Transmission electron microscopy (TEM) and crystallography analysis

Samples for conventional TEM were concentrated from 2-3 mL cultures by centrifugation, adsorbed on carbon-coated copper grids and imaged using a JEOL-1400 Plus (Japan) at 80 kV acceleration. High-resolution (HRTEM) images and selected-area electron diffraction (SAED) patterns were obtained using the TEM mode of a Talos F200X G2 instrument (Thermo Fisher Scientific, Waltham, MA, USA) at 200 kV accelerating voltage. The same device was used for scanning transmission electron microscopy (STEM) high-angle annular dark-field (HAADF) images that were collected for both imaging and mapping elemental compositions by coupling the STEM mode with energy-dispersive X-ray spectrometry (EDS).

### Comparative genomics

Comparative analysis was implemented using complete and near complete genome sequences (Supplementary Table S3). Orthologous protein groups in the transgenic strains with the integrated magnetosome cassette were identified using the OrthoMCL clustering algorithm implemented in *get_homologues* software^63^ with the following parameters: the cut-off threshold for pairwise *blastp* alignments *E value* < 10^-5^ and a minimal protein coverage of 50%. The nonredundant dataset comprising the candidate auxiliary genes for magnetosome formation was constructed by combining the genes identified by transposon mutagenesis in Silva, Schüler et al. (2020) with the genes in the previous experimental studies (Supplementary Table S4). The pangenome matrix generated by *get_homologues* and the script used to compare the orthologues are available under the link: https://github.com/MarDZiuba/pangenomemtx_to_Venn.git

### Graphics and data visualization

All plots were built using in-house scripts written in Python v. 3.8 using *seaborn 0.11.2* ^64^ and *matplotlib* libraries. The magnetosome number and diameters were measured manually using imageJ v.1.53c. The statistical tests and the visualization of their results on the graphs were conducted using the python library *statannot* (https://github.com/webermarcolivier/statannot).

## Data availability statement

Sequencing reads originating from this study were deposited to NCBI GenBank under the BioProject number PRJNA923495.

## Acknowledgements

This study was supported by the European Research Council (ERC) under the European Union’s Horizon 2020 research and innovation program (Grant No. 692637 to D.S.). We are grateful to M. Schüler and S. Geimer for their help with electron microscopy. Electron microscopy performed at University of Pannonia was supported by the National Research, Development and Innovation Office (Hungary) (Grant No. RRF-2.3.1-21-2022-00014). We also thank A. Hübner, L. Borgert, J. Kachel, and B. Melzer for technical assistance.

---

**Extended Data Fig. 1.**
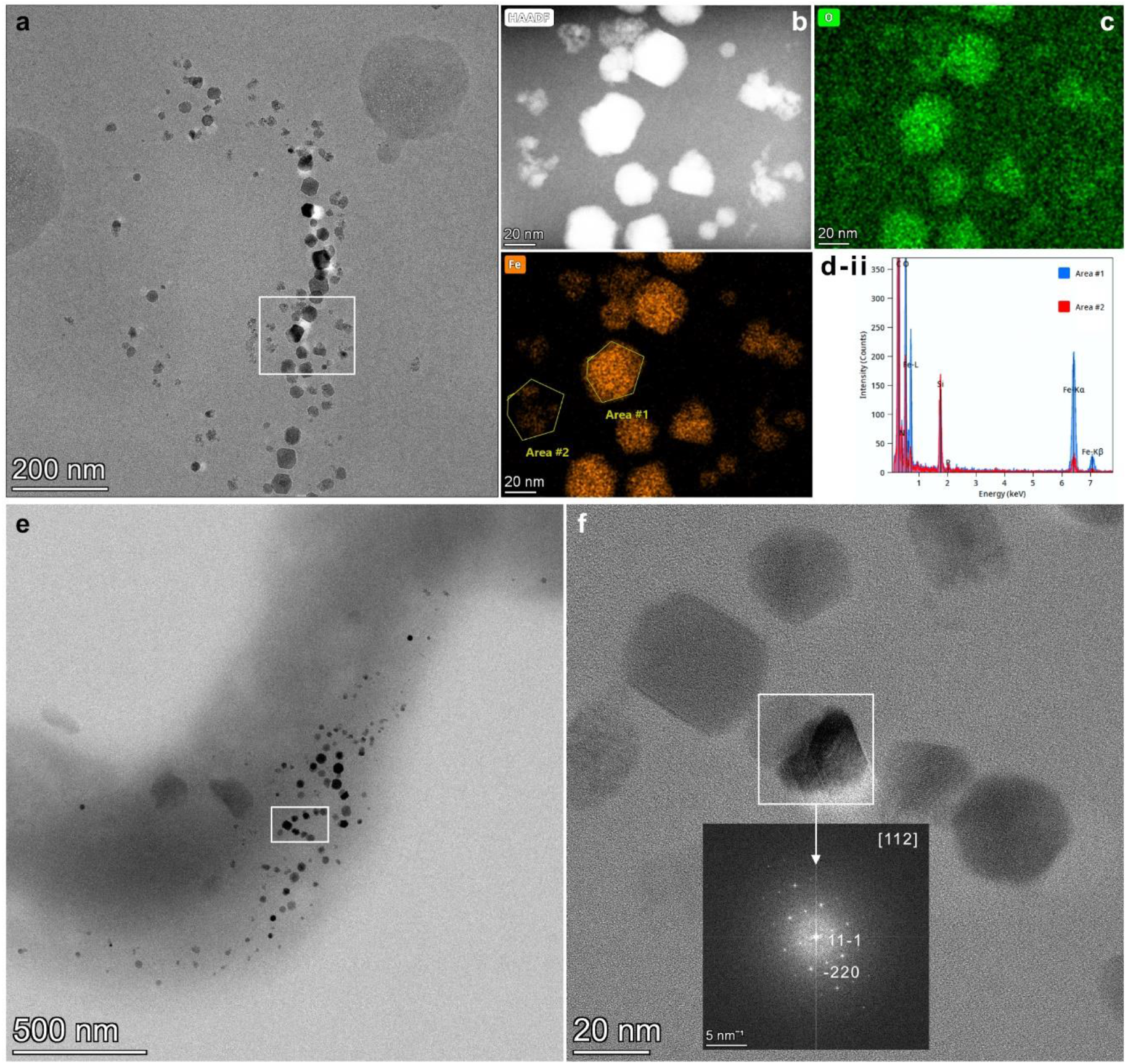
Crystallographic analysis of the magnetosomes produced by B. viridis MAG. (a) and (e) BF TEM micrographs of the magnetosome chain. HAADF (b) and elemental maps (c – d-i) of the boxed area in (a). (d-ii) EDS spectra of the areas marked in (d-i). (f) HRTEM image of the magnetosome chain from the boxed are in (e) with FFT of the twinned particle in the center in [112] zone-axis orientation of magnetite.

**Extended Data Fig. 2.**
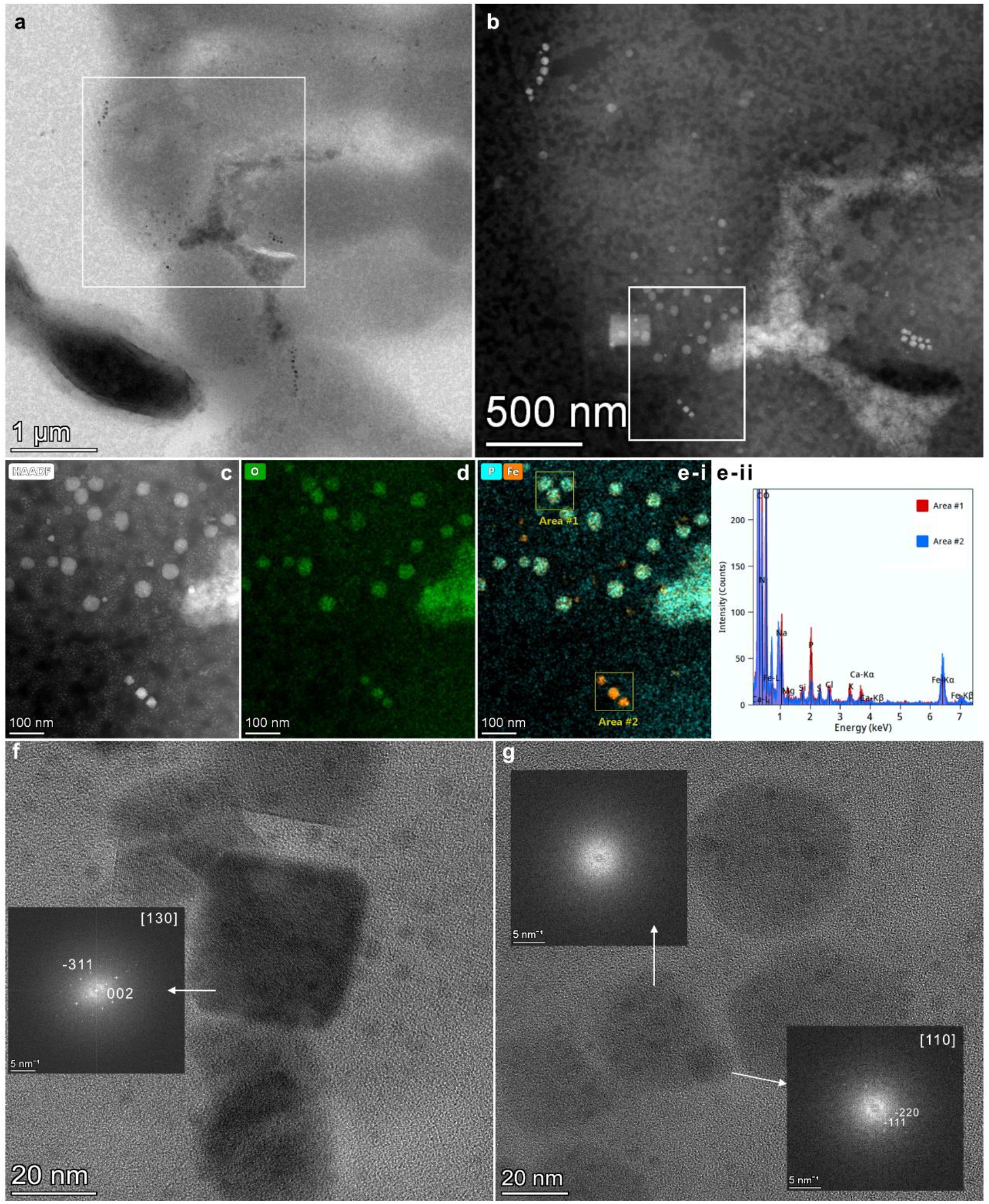
Crystallographic analysis of the magnetosomes produced by Rbl. acidophilus MAG. (a) BF TEM micrograph of the cells. (b) HAADF of the boxed area in (a) with (c-e) close-up HAADF and elemental maps of the close-up fragment in the box in (b). (e-ii) EDS spectra of the areas marked in (e-i). Fe-O-P-rich particles in area #1 likely represent ferrosomes, whereas particles in area #2 are magnetite magnetosomes. (f) and (g) with FFT of the indicated particles. The particle in (f) and the lower particle in (g) are magnetite in [130] and [110] zone-axis orientations, respectively, whereas the upper particle in (g) is amorphous (a putative ferrosome).

**Extended Data Fig. 3.**
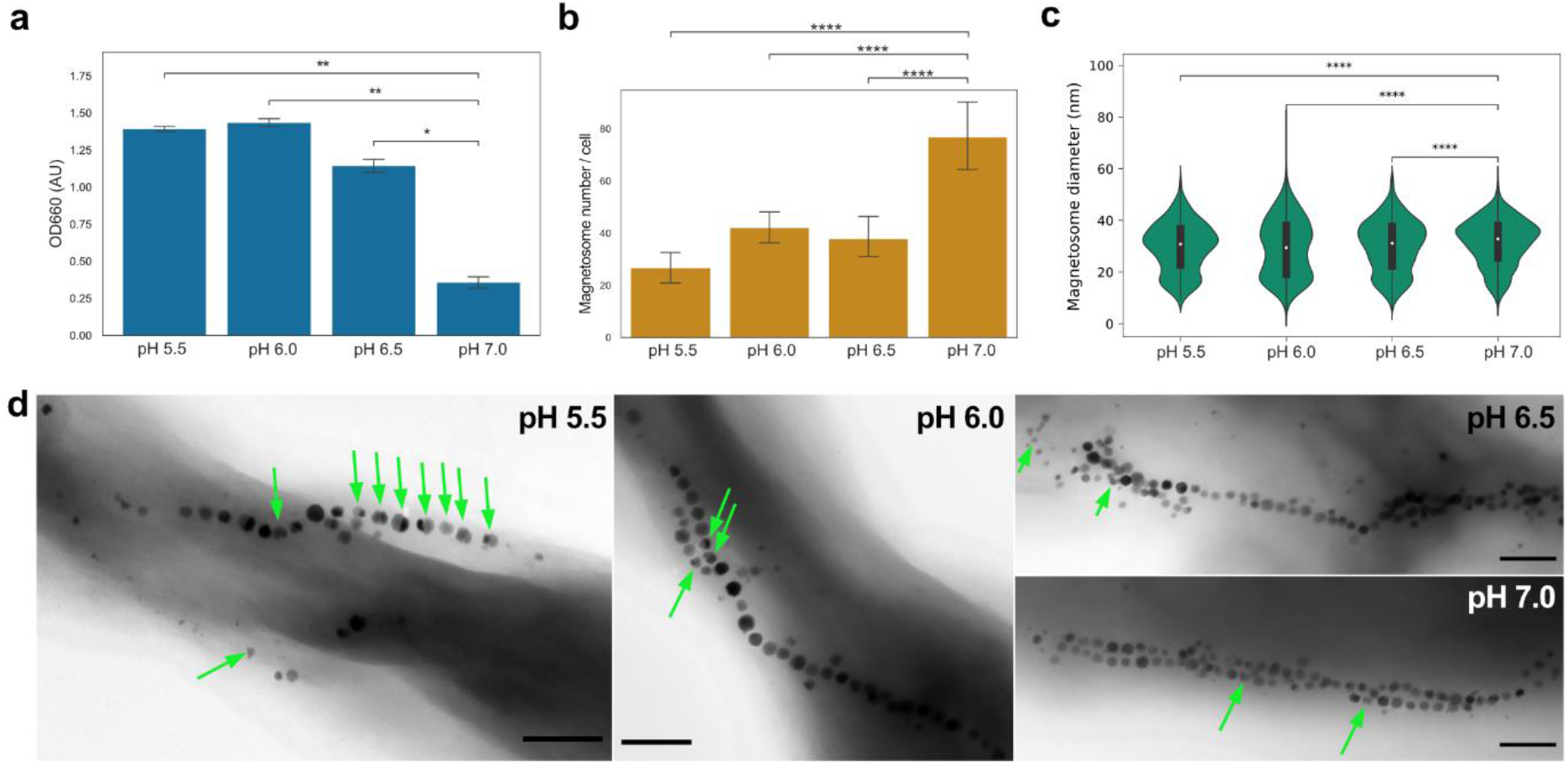
Impact of pH on the magnetosome biosynthesis in Rbl. acidophilus MAG. (a) Bar chart demonstrating the maximum optical density reached by Rbl. acidophilus MAG at different pH. Bars indicate standard deviation. (b) Bar chart demonstrating the average magnetosome number formed per cell at different pH. Bars indicate standard deviation. Number of cells measured ≥;30. (c) Violin plots of the magnetosome diameters at different pH. At least 795 particles from 30 individual cells were measured. (d) TEM micrographs of the magnetosome chains formed at different pH. Arrows indicate defect magnetosome crystals. Scale bars: 200 nm. Statistical significance in (b) was estimated using t-test, and in (c - d) Kruskal-Wallis test, with the subsequence pairwise comparison using Mann-Whitney U test. * Indicate p value < 0.05, ** 0.001 < p value < 0.01, *** 0.0001 < p value < 0.001, **** p value < 0.0001.

**Extended Data Fig. 4.**
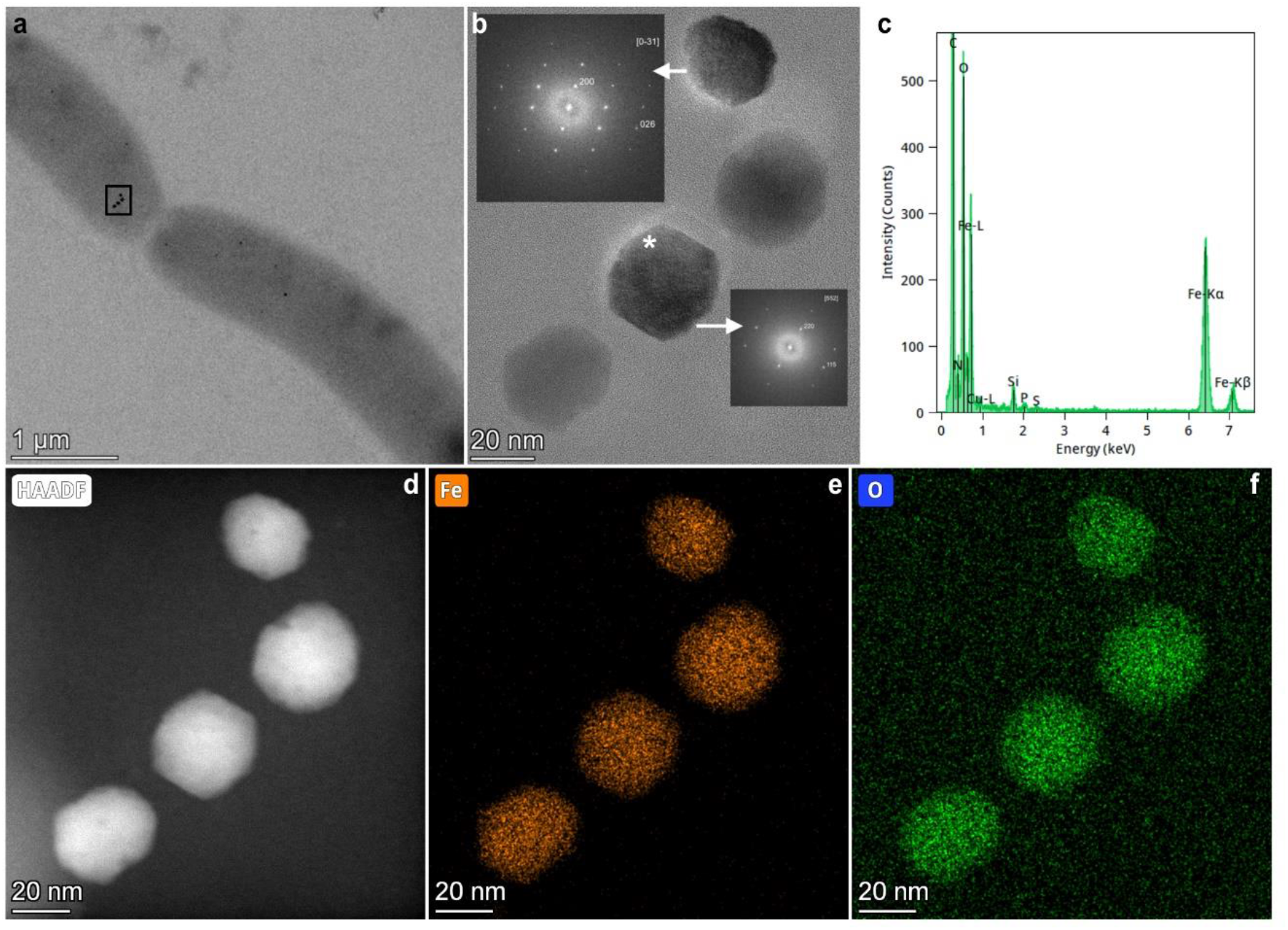
*Crystallographic analysis of the magnetosomes produced by A. brasilense* MAG. (a) BF TEM micrograph of the cells with magnetosomes. (b) HAADF of the magnetosomes from the boxed area in (a) with FFTs of the magnetosomes in zone-axis orientations consistent with magnetite. (c) *EDS spectrum of the magnetosome marked with an asterisk in (b). (d-f) HAADF and elemental maps of the magnetosome chain*.

