## Supplementary material for "Exploring the host range for genetic transfer of magnetic organelle biosynthesis": Table S1

**Supplementary Table S1** Strains used in the study and the experiment outcome

| **N** | **Species Name** | **Strain** | **NCBI Taxonomy: class; order; family** | **Magnetosome Gene Transfer** | **Confirmed Integrity of the inserted gene clusters** | **Magnetosome Biosynthesis** | **Magnetosome Size, nm: Mean Value ± St. Dev.** | **Magnetosome Number/Cell, Mean Value ± St. Dev.** | **Magnetosome Shape** |
| --- | --- | --- | --- | --- | --- | --- | --- | --- | --- |
| 1. | Azospirillum brasilense | Sp7 | Alphaproteobacteria; Rhodospirillales; Azospirillaceae | Yes | Yes | Yes | 24.98 ± 8.56 | 7.56 ± 4.82 | Normal |
| 2. | Thalassospira lucentensis | DSM 14000 | Alphaproteobacteria; Rhodospirillales; Thalassospiraceae | No | NA | - | - | - | - |
| 3. | Bradyrhizobium japonicum | DSM 30131 | Alphaproteobacteria; Hyphomicrobiales; Nitrobacteraceae | No | NA | - | - | - | - |
| 4. | Agrobacterium tumefaciens | C58 | Alphaproteobacteria; Hyphomicrobiales; Rhizobiaceae | Yes | Yes | No | - | - | - |
| 5. | Mesorhizobium japonicum | MAFF303099 | Alphaproteobacteria; Hyphomicrobiales; Phyllobacteriaceae | Yes | Yes | No | - | - | - |
| 6. | Cereibacter sphaeroides | 2.4.1 | Alphaproteobacteria; Rhodobacterales; Rhodobacteraceae | Yes | Yes | Yes | 17.70 ± 6.63 | 5.38 ± 2.94 | Most crystals are twinned or tripled |
| 7. | Rhodobacter capsulatus | SB1003 | Alphaproteobacteria; Rhodobacterales; Rhodobacteraceae | Yes | Yes | No | - | - | - |
| 8. | Rhodobacter capsulatus | B10S | Alphaproteobacteria; Rhodobacterales; Rhodobacteraceae | Yes | Yes | No | - | - | - |
| 9. | Rhodocista centenaria | DSM 9894 | Alphaproteobacteria; Rhodospirillales; Rhodospirillaceae | Yes | No* | No | - | - | - |
| 10. | Pararhodospirillum photometricum | DSM 122 | Alphaproteobacteria; Rhodospirillales; Rhodospirillaceae | No | NA | - | - | - | - |
| 11. | Telmatospirillum siberiense | DSM 18240 | Alphaproteobacteria; Rhodospirillales; Rhodospirillaceae | No | NA | - | - | - | - |
| 12. | Magnetospirillum molischianum | DSM 120 | Alphaproteobacteria; Rhodospirillales; Rhodospirillaceae | No | NA | - | - | - | - |
| 13. | Magnetospirillum fulvum | DSM 113 | Alphaproteobacteria; Rhodospirillales; Rhodospirillaceae | No | NA | - | - | - | - |
| 14. | Rhodomicrobium vannielii | DSM 166 | Alphaproteobacteria; Hyphomicrobiales; Hyphomicrobiaceae | Yes | Yes | Yes | 30.44 ± 9.95 | 33.73 ± 31.25 | Normal |
| 15. | Rhodopseudomonas pseudopalustris | DSM 123 | Alphaproteobacteria; Hyphomicrobiales; Nitrobacteraceae | Yes | Yes | Yes | 17.81 ± 6.97 | 42.23 ± 27.67 | Most crystals are flake-shaped |
| 16. | Rhodovibrio salinarum | DSM 9154 | Alphaproteobacteria; Hyphomicrobiales; Rhodovibrionaceae | No | NA | - | - | - | - |
| 17. | Rhodoblastus acidophilus | DSM 137 | Alphaproteobacteria; Hyphomicrobiales; Beijerinckiaceae | Yes | Yes | Yes | 35.93 ± 9.42 | 33.76 ± 28.40 | Normal at pH 6.5-7.0 |
| 18. | Rhodoplanes elegans | DSM 11907 | Alphaproteobacteria; Hyphomicrobiales; Hyphomicrobiaceae | Yes | Yes | Yes | 19.19 ± 5.33 | 3.80 ± 2.59 | Normal |
| 19. | Blastochloris viridis | DSM 133 | Alphaproteobacteria; Hyphomicrobiales; Blastochloridaceae | Yes | Yes | Yes | 24.60 ± 9.29 | 46.43 ± 39.70 | Normal, numerous flake-shaped crystals are also present |
| 20. | Rhodobium orientis | DSM 11290 | Alphaproteobacteria; Hyphomicrobiales; Rhodobiaceae | No | NA | - | - | - | - |
| 21. | Rhodothalassium salexigens | DSM 2132 | Alphaproteobacteria; Rhodothalassiales; Rhodothalassiaceae | No | NA | - | - | - | - |
| 22. | Rubrivivax gelatinosus | DSM 1709 | Betaproteobacteria; Burkholderiales; Burkholderiales genera incertae sedis | Yes | Yes | No | - | - | - |
| 23. | Rhodocyclus tenuis | DSM 109 | Betaproteobacteria; Rhodocyclales; Rhodocyclaceae | No | NA | - | - | - | - |
| 24. | Allochromatium vinosum | DSM 180 | Gammaproteobacteria; Chromatiales; Chromatiaceae | No | NA | - | - | - | - |
| 25. | Wolinella succinogenes | DSM1740 | Epsilonproteobacteria; Campylobacterales; Helicobacteraceae | No | NA | - | - | - | - |

NA not applicable

* Large deletion affecting most of the inserted cassette
