## Supplementary material for "Exploring the host range for genetic transfer of magnetic organelle biosynthesis": Table S2

**Supplementary Table S2 Cultivation conditions**

| **Species Name** | **Strain** | **Medium used for cultivation (with references)** | **Kanamycin**  **concentration for transconjugants selection, µg/mL** | **Growth Temperature,** °C | **Cultivation conditions** | | | **Source** |
| --- | --- | --- | --- | --- | --- | --- | --- | --- |
|  |  |  |  |  | **Routine cultivation** | **Agar plates** | **For potential magnetosome formation** |  |
| *Azospirillum brasilense* | Sp7 | Flask standard medium (FSM) ^1^ | 25 | 28 | Aerobic, in the dark | Aerobic, in the dark | Microoxic or anaerobic, in the dark | Lab collection |
| *Thalassospira lucentensis* | DSM 14000 | Bacto Marine Broth (DIFCO 2216) | 40 | 18 | Aerobic, in the dark | Aerobic, in the dark | NA* | DSMZ** |
| *Bradyrhizobium japonicum* | DSM 30131 | Flask standard medium (FSM) ^1^ or PY (Pro 1L: 3 g soybean peptone, 3 g yeast extract) | 150 | 28 | Aerobic, in the dark | Aerobic, in the dark | NA | Lab collection |
| *Agrobacterium tumefaciens* | C58 | Flask standard medium (FSM) ^1^ or PY (Pro 1L: 3 g soybean peptone, 3 g yeast extract) | 50 | 28 | Aerobic, in the dark | Aerobic, in the dark | Microoxic or anaerobic, in the dark | Lab collection |
| *Mesorhizobium japonicum* | MAFF303099 | Flask standard medium (FSM) ^1^ or PY (Pro 1L: 3 g soybean peptone, 3 g yeast extract) | 150 | 28 | Aerobic, in the dark | Aerobic, in the dark | Microoxic or anaerobic, in the dark | Lab collection |
| *Cereibacter sphaeroides* | 2.4.1 | Sistrom medium ^2^ or FSM^1^ | 10 | 28 | Anaerobic or microoxic, with light | Aerobic, with light | Anaerobic, with light | DSMZ |
| *Rhodobacter capsulatus* | SB1003 | Sistrom medium ^2^ or FSM^1^ | 10 | 28 | Anaerobic or microoxic, with light | Aerobic, with light | Anaerobic, with light | A kind gift from T. Drepper (Research Center Jülich, Germany) |
| *Rhodobacter capsulatus* | B10S | Sistrom medium ^2^ or FSM^1^ | 10 | 28 | Anaerobic or microoxic, with light | Aerobic, with light | Anaerobic, with light | A kind gift from T. Drepper (Research Center Jülich, Germany) |
| *Rhodocista centenaria* | DSM 9894 | CENMED^3^ | Because of the native resistance of the strain to kanamycin, a version of pTpsMAG1 with spectinomycin (Sp) resistance was used for transfers. Sp was applied at 10 µg/ml. | 37 | Anaerobic or microoxic, with light | Aerobic, with light | NA | DSMZ |
| *Pararhodospirillum photometricum* | DSM 122 | Rhodospirillaceae medium 27, as described on DSMZ webpage (<https://www.dsmz.de/microorganisms/medium/pdf/DSMZ_Medium27.pdf>) | 5 | 28 | Anaerobic, with light | Anaerobic, with light | NA | DSMZ |
| *Telmatospirillum siberiense* | DSM18240 | FSM^1^ | 5 | 28 | Anaerobic, in the dark | Anaerobic, in the dark | NA | DSMZ |
| *Magnetospirillum molischianum* | DSM120 | Rhodospirillaceae medium 27, as described on DSMZ webpage (<https://www.dsmz.de/microorganisms/medium/pdf/DSMZ_Medium27.pdf>) | 10 | 28 | Anaerobic, with light | Anaerobic, with light | NA | DSMZ |
| *Magnetospirillum fulvum* | DSM113 | Rhodospirillaceae medium 27, as described on DSMZ webpage (<https://www.dsmz.de/microorganisms/medium/pdf/DSMZ_Medium27.pdf>) | 10 | 28 | Anaerobic, with light | Anaerobic, with light | NA | DSMZ |
| *Rhodomicrobium vannielii* | DSM 166 | FSM^1^ | 0.5 | 28 | Anaerobic or microoxic, with light | Aerobic, with light | Anaerobic, with light | Lab collection |
| *Rhodopseudomonas pseudopalustris* | DSM123 | YPS (Pro 1L: 3 g soybean peptone, 3 g yeast extract, 2 mM MgSO_4_, 2 mM CaCl_2_) for conjugations, FSM^1^ for magnetosome formation | 100 | 28 | Anaerobic or microoxic, with light | Aerobic, with light | Anaerobic, with light | Lab collection |
| *Rhodovibrio salinarum* | DSM 9154 | Halophile Medium 652 (<https://www.dsmz.de/microorganisms/medium/pdf/DSMZ_Medium652.pdf>) or FSM^1^ supplemented with 10% NaCl | 25 | 37 | Anaerobic or microoxic, with light | Aerobic, with light | NA | DSMZ |
| *Rhodoblastus acidophilus* | DSM 137 | Acid *Rhodospirillaceae* Medium 26 (pH 5.5) as described on DSMZ webpage (<https://www.dsmz.de/microorganisms/medium/pdf/DSMZ_Medium26.pdf>), FSM-MES (FSM^1^ with 10 mM MES buffer instead of HEPES, pH 5.5-7.0) | 10 | 28 | Anaerobic with light | Anaerobic with light | Anaerobic, with light | DSMZ |
| *Rhodoplanes elegans* | DSM 11907 | FSM-AmPyr (FSM^1^ with sodium nitrate substituted with 4 mM ammonium chloride and potassium lactate substituted with 15 mM sodium pyruvate) | 30 | 28 | Anaerobic with light | Anaerobic with light | Anaerobic, with light | DSMZ |
| *Blastochloris viridis* | DSM 133 | FSM-AmPyr (FSM^1^ with sodium nitrate substituted with 4 mM ammonium chloride and potassium lactate substituted with 15 mM sodium pyruvate) | 20 | 28 | Anaerobic with light | Anaerobic with light | Anaerobic, with light | DSMZ |
| *Rhodobium orientis* | DSM 11290 | Rhodobium medium 745, as described on DSMZ webpage (<https://www.dsmz.de/microorganisms/medium/pdf/DSMZ_Medium745.pdf>) | 10 | 28 | Anaerobic with light | Anaerobic with light | NA | DSMZ |
| *Rhodothalassium salexigens* | DSM 2132 | YPS (Pro 1L: 3 g soybean peptone, 3 g yeast extract, 2 mM MgSO_4_, 2 mM CaCl_2_) or FSM^1^ supplemented with 3-5% NaCL | 25 | 28 | Anaerobic with light | Anaerobic with light |  | DSMZ |
| *Rubrivivax gelatinosus* | DSM 1709 | FSM^1^ | 50 | 28 | Anaerobic or microoxic, with light | Aerobic, with light | Anaerobic, with light | DSMZ |
| *Rhodocyclus tenuis* | DSM 109 | Rhodospirillaceae medium 27, as described on DSMZ webpage (<https://www.dsmz.de/microorganisms/medium/pdf/DSMZ_Medium27.pdf>) | 5 | 28 | Anaerobic with light | Anaerobic with light | NA | DSMZ |
| *Allochromatium vinosum* | DSM 180 | MYE^4^ | 10 | 28 | Anaerobic with light | Anaerobic with light | NA | DSMZ |
| *Escherichia coli* | WM3064 | LB (lysogeny broth) supplemented with 0.1 mM DL-α,ε-diaminopimelic acid (DAP) | 25 | 37 | Aerobic, in the dark | Aerobic, in the dark | Microoxic or anaerobic, in the dark | Lab collection, William Metcalf, UIUC, unpublished |
| *Wolinella succinogenes* | DSM1740 | Fumarate medium^5^ supplementer with 0.5% (w/v) Brain-Heart-Infusion extract | 25 | 37 | Anaerobic, in the dark | Anaerobic, in the dark | NA | A kind gift from J. Simon (Centre for Synthetic Biology, Technical University of Darmstadt, Germany) |

*Not applicable, i.e., no mutants for magnetosome formation were obtained

** DSMZ-German Collection of Microorganisms and Cell Cultures GmbH
