## Supplementary material for "Exploring the host range for genetic transfer of magnetic organelle biosynthesis": Table S3

**Supplementary Table S3 Genomes used for the orthologues search and comparative analysis**

| **Strain** | **Genome Assembly** | **Assembly Level** | **CheckM**^1^ **completeness*** | **CheckM contamination (marker duplication)*** |
| --- | --- | --- | --- | --- |
| Magnetospirillum gryphiswaldense MSR-1 | [CP027526.1](https://www.ncbi.nlm.nih.gov/nuccore/CP027526.1/) | Complete | 99.9% | 0% |
| Rhodospirillum rubrum ATCC 11170 | [GCA_000013085.1](https://www.ncbi.nlm.nih.gov/assembly/GCA_000013085.1) | Complete | 99.5% | 0% |
| Magnetospirillum sp. 15-1 | GCA_900184795.1 | Contig | 98.5% | 3.5% |
| Rhodovastum atsumiense G2-11 | [GCA_937425535.1](https://www.ncbi.nlm.nih.gov/assembly/GCA_937425535.1) | Complete | 99.5% | 0.5% |
| Cereibacter sphaeroides 2.4.1 | [GCA_000012905.2](https://www.ncbi.nlm.nih.gov/assembly/GCA_000012905.2) | Complete | 99.4% | 0% |
| Rhodobacter capsulatus SB 1003 | GCA_000021865.1 | Complete | 99.1% | 0.6% |
| Rhodomicrobium vannielii ATCC 17100 | GCA_000166055.1 | Complete | 97.8% | 0.3% |
| Mesorhizobium japonicum MAFF 303099 | GCA_000009625.1 | Complete | 99.9% | 2.0% |
| Blastochloris viridis DSM 133 | GCA_001548155.2 | Complete | NA | NA |
| Rhodoblastus acidophilus DSM 137 | GCA_900187365.1 | Contig | 98.4% | 1.7% |
| Rhodopseudomonas pseudopalustris DSM 123 | GCA_900110435.1 | Scaffold | NA | NA |
| Rhodoplanes elegans DSM 11907 | GCA_003258805.1 | Scaffold | NA | NA |
| Agrobacterium fabrum str. C58 | GCA_000092025.1 | Complete | 100.0% | 0.0% |
| Azospirillum brasilense Sp7 | GCA_001315015.1 | Complete | 100.0% | 3.3% |
| Rubrivivax gelatinosus DSM 1709 | GCA_004340905.1 | Scaffold | 99.8% | 0.5% |

*From <https://mage.genoscope.cns.fr/>
