## Supplementary material for "Exploring the host range for genetic transfer of magnetic organelle biosynthesis": Fig. S

Supplementary Figures for the manuscript.

### Table of content:

Supplementary methods

Figure S1

Figure S2

Figure S3

Figure S4

### Supplementary methods

Figure S1: A multiple alignment of full length 16S rRNA genes was performed with MAFFT v7. using default settings, and highly variable regions that produced ambiguous alignments were manually trimmed. Phylogenetic analysis was conducted using The maximum likelihood trees were computed using *iqtree* v. 2.1.3[1] with automatic model selection by ModelFinder[2], performing 1000 ultrafast bootstrap (UFBoot)[3] replicates.

Figure S2: The fragments were amplified using Paq5000 dna polymerase with the primer sets published previously[4]. The fragments were separated in 1% agarose gels with TAE buffer as described elsewhere[5].

Figure S3: TEM grids preparation as described in the main text.

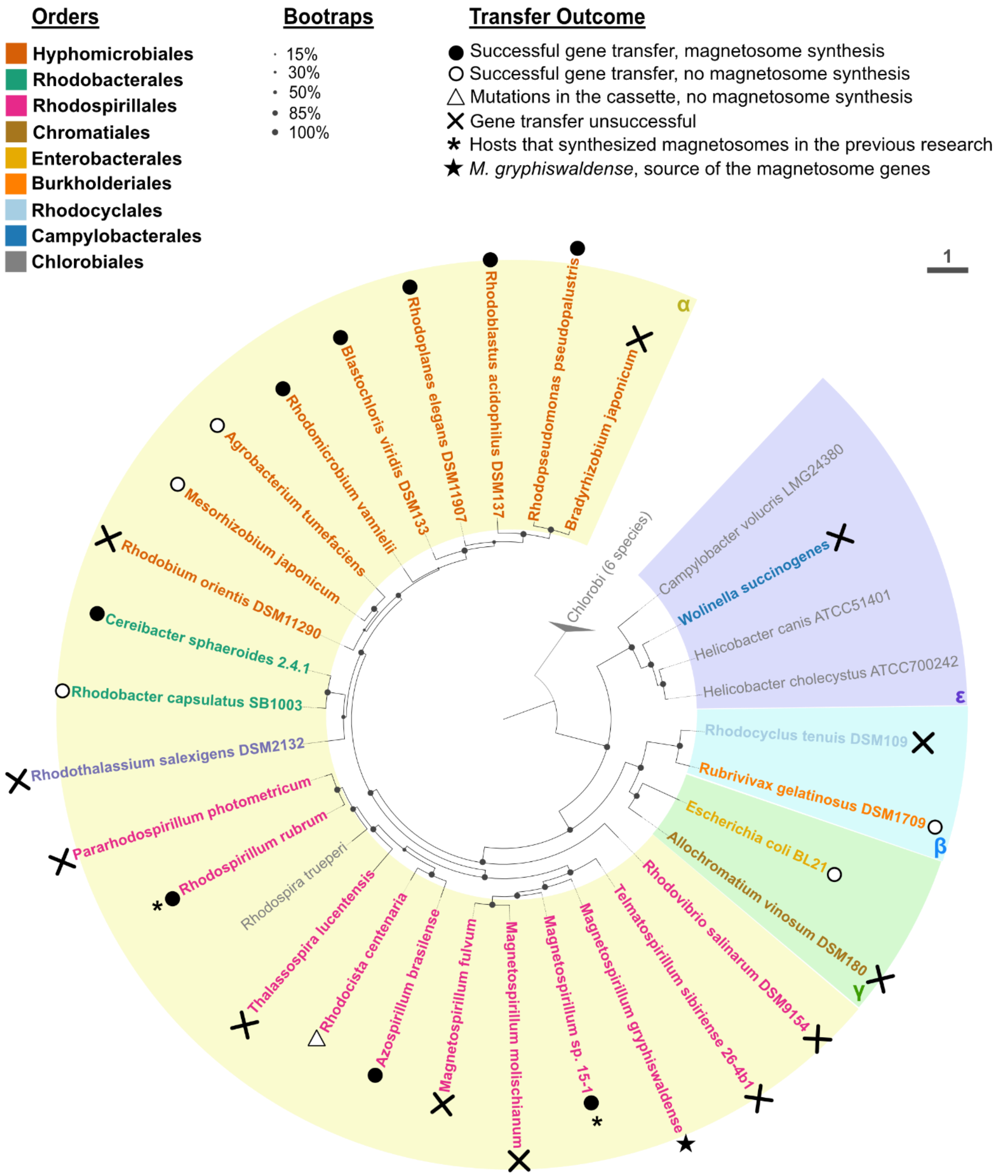


Figure S1 Phylogenetic tree based on 16S rRNA gene (maximum likelihood) illustrating the relationships between the strains used in the study. Circles at nodes indicate branch support calculated from 1000 replicates using ultrafast bootstrap analysis. Representatives of *Chlorobi* were used as an outgroup. Branch length represents the number of base substitutions per site. Strains indicated in bold were used as recipients for the transfer of the magnetosome genes. The colored shadings highlight classes: "α, β, γ, ε" correspond to *Alpha*-, *Beta*-, *Gamma*- and *Epsilonproteobacteria*. The transfer experiments outcomes are indicated by symbols.

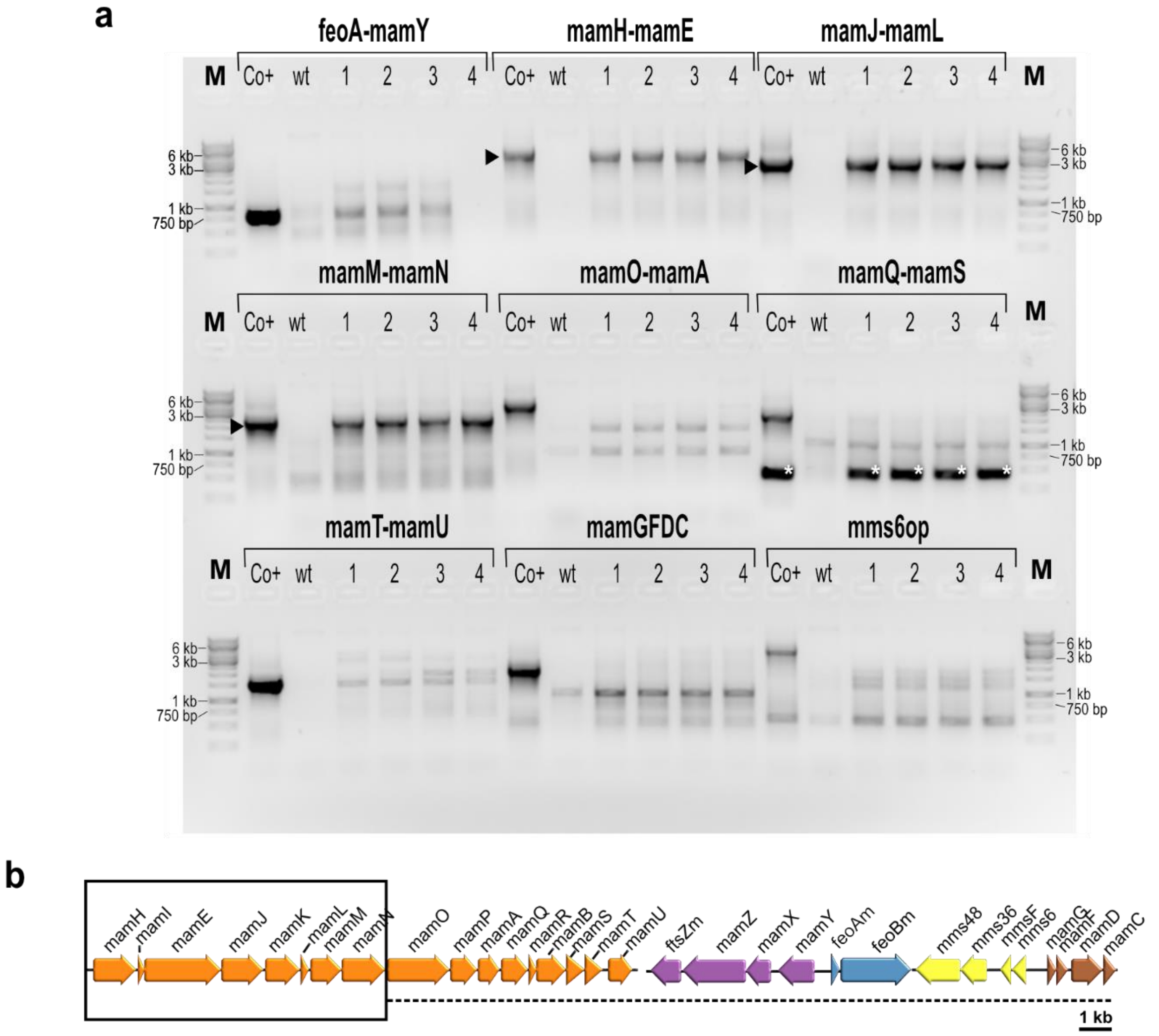


Figure S2 PCR-analysis of the magnetosome cassette in *Rh. centenaria* MAG mutants. (a) Agarose gel of the PCR fragments amplified from different regions of the MAG cassette. "M": marker; "Co+": plasmid pTpsMAG1 as a positive control; "wt": wildtype strain of *Rh. centenaria;* "1-4" randomly selected transconjugants of *Rh. centenaria* MAG. Arrowheads indicate the correct bands, asterisks in mamQ-mamS region indicate a non-specific band corresponding to a fragmen within gene *mamE*. (b) Schematic illustrating the mutation occurred in *Rh. centenaria* MAG mutants as determined using the PCR reactions shown in (a): the region in a rectangle is preserved, whereas the fragment underlined with the punctuated line is deleted.

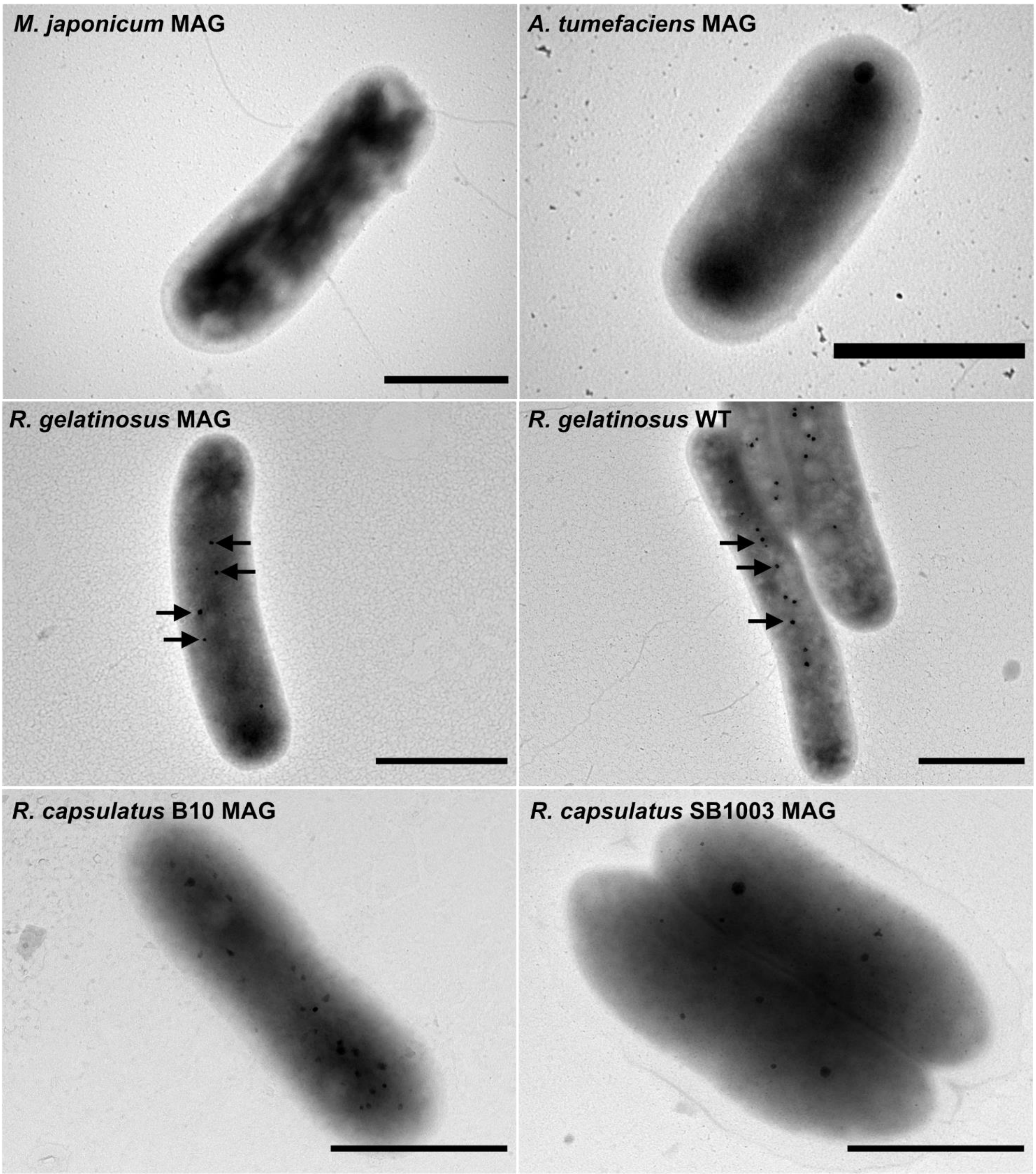


Figure S3 Electron micrographs of the MAG mutants that did not synthesize magnetosomes. Note that the conspicuous electron-dense particles iindicated by arrows in *Rubrivivax gelatinosus* are also present in the wildtype and, hence, are not related to magnetosome fromation. Scale bars: 1 µm.

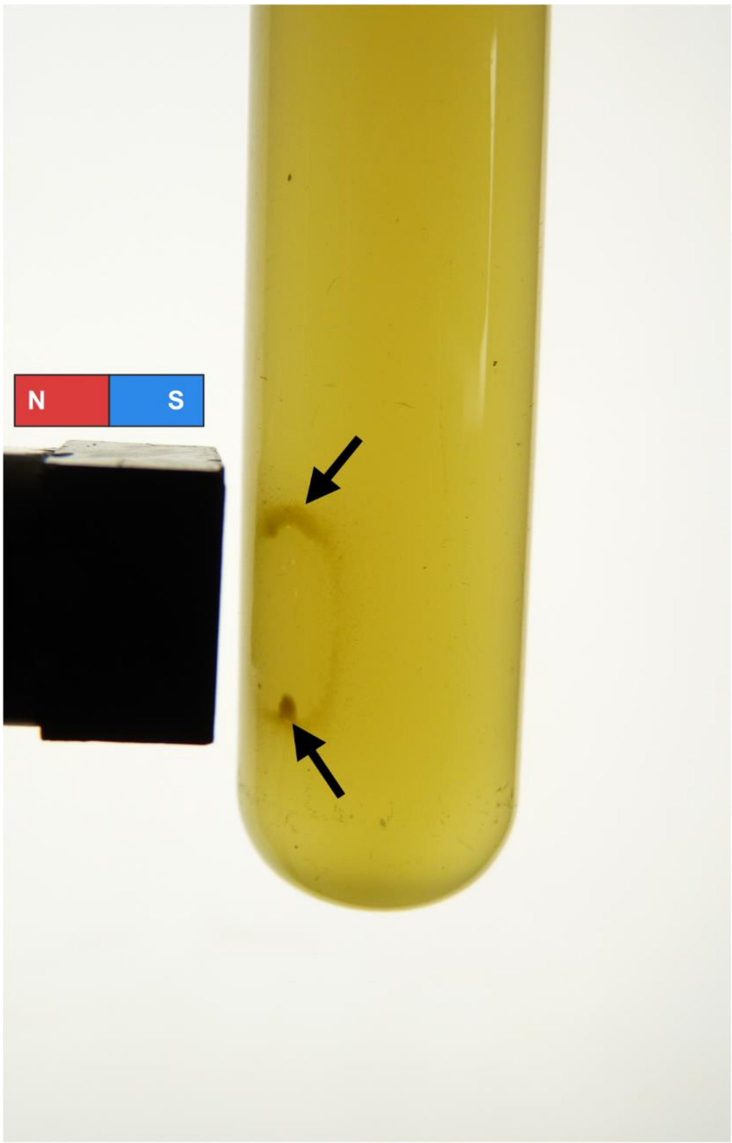


Figure S4 Culture of *B. viridis* MAG with cells accumulated near the magnet pole (indicated by arrows).
